# The combination of nelfinavir and cisplatin drives lytic cell death through a caspase-8/caspase-3/GSDME axis in platinum-resistant ovarian cancer cells

**DOI:** 10.64898/2026.06.30.735544

**Authors:** Benjamin Forgie, Rewati Prakash, Desiree Marno, Farah H. Abdalbari, Edith Zorychta, Abu Shadat M. Noman, Alicia A. Goyeneche, Lucy Gilbert, Julia V. Burnier, Carlos M. Telleria

**Affiliations:** Cancer Research Program, Research Institute of the McGill University Health Centre, Montreal, Quebec, Canada; Department of Pathology, McGill University, Montreal, Quebec, Canada; Department of Pharmacology and Therapeutics, McGill University, Montreal, Quebec, Canada; Department of Biochemistry and Molecular Biology, Chittagong University, Chittagong, Bangladesh; Division of Gynecologic Oncology, Research Institute of the McGill University Health Centre, Montreal, Quebec, Canada; Gerald Bronfman Department of Oncology, McGill University, Montreal, Quebec, Canada

**Author notes:** Correspondence: Carlos M. Telleria.

**Keywords:** Ovarian cancer, Cisplatin, Nelfinavir, Pyroptosis, Gasdermin E, Caspase-8, Endoplasmic reticulum stress, Drug resistance

## Abstract

**Purpose:** Cisplatin (CDDP) is the most active chemotherapy for ovarian cancer; primary or acquired resistance signals a poor prognosis. Nelfinavir (NFV), an HIV protease inhibitor, has demonstrated anti-tumor activity in multiple cancer models, but its interaction with CDDP in ovarian cancer has yet to be demonstrated. In this work, we addressed whether the combination of CDDP and NFV provides treatment advantage in platinum (Pt)-resistant ovarian cancer cells.

**Methods:** Drug synergy between NFV and CDDP was assessed using cell vitality assays and Loewe additivity modelling. Apoptotic and pyroptotic signalling were evaluated by immunoblotting, mitochondrial membrane potential analysis, and lactate dehydrogenase (LDH) release, and caspase inhibition. Transcriptomic changes were assessed by bulk mRNA sequencing followed by differential gene expression analysis and gene set enrichment analysis.

**Results:** NFV synergized with CDDP to reduce the viability of Pt-resistant ovarian cancer cells, promoting a regulated lytic cell death phenotype involving apoptotic and pyroptotic features. Combination treatment induced caspase-8 and caspase-3 activation, and downstream gasdermin E (GSDME) processing. Inhibition of caspase-3 significantly attenuated cell death, and caspase-8 inhibition rescued viability and prevented Bid cleavage, caspase-3 activation, and GSDME cleavage. These effects occurred in the context of enhanced endoplasmic reticulum stress, increased DNA damage with reduced DNA repair, and impaired Akt-driven survival signalling.

**Conclusions:** Our findings establish that NFV synergizes with CDDP in killing Pt-resistant ovarian cancer cells by promoting a caspase-8-dependent apoptotic-to-secondary pyroptotic response, supporting further investigation of NFV as a potential drug to be repurposed to increase the efficacy of Pt-based therapy.

## 1. Introduction

Ovarian cancer is the most lethal gynecological malignancy and remains a major clinical challenge due to frequent diagnosis at advanced stage and high rates of disease recurrence [1, 2]. Standard-of-care treatment comprises cytoreductive surgery in combination with platinum (Pt)-based chemotherapy [3]. Despite initial responsiveness to Pt-based chemotherapy, most patients with ovarian cancer ultimately develop recurrent Pt-resistant disease, underscoring the need for strategies that enhance treatment efficacy [4]. Pt resistance arises through a network of adaptive changes that collectively reduce treatment sensitivity and limit cell death; these include mechanisms such as dysregulation of Pt influx and efflux transporters [5], increased DNA damage repair capacity [6], activation of stress-adaptive programs [7, 8], and engagement of pro-survival signalling pathways such as PI3K-Akt while simultaneously weakening pro-death signalling [4, 9, 10]. As a result, Pt-resistant cells may tolerate substantial Pt-induced damage yet fail to convert that stress into efficient cell death [11]. Accordingly, there is considerable interest in identifying agents that can enhance the cytotoxic effects of Pt drugs and overcome these adaptive survival responses in ovarian cancer cells.

Nelfinavir (NFV), a human immunodeficiency virus (HIV) protease inhibitor [12], has emerged as a candidate for drug repurposing in oncology [13] due to its reported anti-tumor activity across multiple cancer types [14]. We have previously demonstrated that NFV exerts cytotoxic effects against high-grade serous ovarian cancer (HGSOC) cells through engagement of pathways linked to the unfolded protein response (UPR), DNA damage, and pro-apoptotic signalling [15]. Additionally, NFV has been shown to inhibit Akt and Erk mediated pro-survival signalling [15]. Together, these findings suggest that NFV may enhance the cytotoxicity of Pt-based drugs by increasing cellular stress, suppressing pro-survival signalling, and lowering the threshold for treatment-induced regulated cell death. However, whether NFV interacts with cisplatin (CDDP), the first-generation Pt agent used worldwide [16, 17], to kill Pt-resistant ovarian cancer cells remains unknown.

Pt drugs such as CDDP primarily exert their anti-cancer effects through the formation of detrimental intra- and inter-strand DNA crosslinks [9, 18], which in turn activate the DNA damage response [19] and culminate in engaging the intrinsic apoptotic pathway and the activation of executioner caspases -3 and -7 [9, 18]. In some contexts, apoptotic signalling may also be reinforced through activation of initiator caspases such as caspase-8 [20], which can link upstream death signalling to mitochondrial apoptotic amplification through cleavage of Bid, a pro-apoptotic BH3-only protein member of the BCL2 family of genes [21], and through direct executioner caspase cleavage [20, 22]. While caspase-3 is canonically associated with apoptotic execution, it can also function as a molecular branch point linking apoptosis to the lytic cell death program known as pyroptosis [23, 24]. Pyroptotic cell death is characterized by the formation of gasdermin (GSDM) protein pores in the plasma membrane, ranging from 10-20 nm in size, permitting ion flux and leakage of small intracellular molecules [25, 26]. Following release of these intracellular materials, terminal plasma membrane rupture is mediated by ninjurin-1 (NINJ1) [27], which oligomerizes in the plasma membrane to form large ring-like openings, ranging 10-100 nm, that enable the leakage of larger molecules such as lactate dehydrogenase (LDH), a marker of lytic cell death [28].In cells expressing sufficient gasdermin E (GSDME), caspase-3 mediated GSDME cleavage liberates the pore-forming N-terminal GSDME fragment, resulting in initial membrane permeabilization followed by NINJ1-mediated cell lysis [23, 24, 29]. Accordingly, the outcome of caspase-3 activation may depend not only on the strength of apoptotic signalling, but also on cellular determinants such as GSDME expression and the upstream signalling environment in which caspase-3 is activated [29]. This suggests that combination therapies capable of amplifying the activity of cell-death inducing caspases may also reshape the modality of regulated cell death, which may involve not only apoptotic, but also pyroptotic molecular mediators.

In this study, we report that the combination of NFV and CDDP synergize in triggering cell death in Pt-resistant ovarian cancer cells and promotes a regulated lytic cell death phenotype that involves caspase-3-mediated GSDME activation. The combination treatment was associated with features of both apoptosis and pyroptosis, suggesting that NFV may alter not only the extent but also the modality of CDDP-induced cell death. We further identified caspase-8 as an upstream mediator of this response, acting prior to Bid cleavage and caspase-3 activation, in a cellular context marked by enhanced ER stress, DNA damage and reduce repair, and impaired survival/growth signalling. Collectively, these findings provide mechanistic insight into how NFV and CDDP potentiate one another in killing Pt-resistant ovarian cancer cells and highlight the translational potential of repurposing NFV as an adjuvant to Pt-based therapy in ovarian cancer.

## 2. Materials and methods

### 2.1 Cell Culture and Reagents

The PEO4, IGROV-1/CP, and TOV21G cell lines were used in this study as models of Pt-resistant ovarian cancer. PEO4 cells were isolated from the ascites of a patient 10 months following their second round of Pt-based chemotherapy and are considered clinically resistant to Pt [30]. IGROV-1/CP cells were derived from the IGROV-1 parental line and were rendered Pt-resistant through chronic exposure to increasing concentrations of CDDP [31]. TOV21G cells were isolated from a patient prior to Pt-exposure [32] and have been shown to be intrinsically Pt-resistant [33]. The PEO4 and TOV21G cell lines were kindly gifted by Dr. Taniguchi (Fred Hutchinson Cancer Centre, University of Washington, WA, USA) and Dr. Mes-Masson (Centre de recherche du Centre Hospitalier de l’Université de Montréal (CHUM), Montréal, QC, Canada), respectively. IGROV-1/CP was obtained from Dr. Stephen Howell (University of California, San Diego, CA, USA). PEO4 cells harbor a homozygous inactivating *TP53* mutation [34], whereas IGROV-1/CP cells carry heterozygous inactivating *TP53* mutations inherited from the parental line [34, 35], and TOV21G cells are wild-type for *TP53* [34]. All cells were cultured in RPMI 1640 media (Mediatech, Manassas, VA, USA) supplemented with 10% fetal bovine serum (FBS) (Corning Inc, Corning, NY, USA), 0.01 mg/mL of human insulin (Roche, Indianapolis, IN, USA), 10 mM HEPES (Corning), 100 IU penicillin (Mediatech), 100 µg/mL streptomycin (Mediatech), 2 mM L-Alanyl-L-glutamine (Glutagro^TM^, Corning), and 1mM sodium pyruvate (Corning). Cells were maintained at 37 °C in a humidified incubator with 5% CO_2_. To complete this study, the following drugs were used: nelfinavir mesylate hydrate (NFV) (Sigma Chemical Co., St. Louis, MO, USA), cis-diamminedichloroplatinum II (CDDP) (Sigma), Z-IETD-FMK (Cedarlane Laboratories, Burlington, ON, Canada), Z-DEVD-FMK (Cedarlane), necrostatin-1 (NEC-1) (Cedarlane), and ferrostatin-1 (FER-1) (Cedarlane). CDDP was dissolved in 0.9% sodium chloride (saline) (Sigma). NFV, Z-IETD-FMK, Z-DEVD-FMK, NEC-1 and FER-1 were dissolved in dimethyl sulfoxide (DMSO) (Sigma), with a maximum concentration of 0.1% (v/v) used in cell culture.

### 2.2 Assessment of cell proliferation and viability

Cell proliferation and viability were assessed through microcapillary cytometry using the Guava Muse Cell Analyzer (Cytek Biosciences, Fremont, CA, USA). Cells were seeded in 6-well plates and treated with increasing concentrations of CDDP for 3 hours (h). Thereafter, the CDDP-containing media were removed, and the cells were incubated for an additional 72 h in the presence of either a constant concentration of 20 µM NFV or drug-free media. The cells were then trypsinized, collected, and stained with the Muse Count and Viability Reagent (Cytek) according to the manufacturer’s protocol. In brief, cells were stained simultaneously with a membrane-permeable dye to obtain the total cell count, and a membrane-impermeable dye to determine the number of cells with decreased membrane integrity (non-viable cells). From these counts, the number of viable cells with intact plasma membranes was determined.

### 2.3 Cell vitality measurement and drug-drug interaction assessment

Drug interaction between CDDP and NFV was assessed using the Loewe additivity model in SynergyFinder [36, 37]. In this model, scores less than -10 indicate antagonism between two drugs, whereas scores from -10 to 10 suggest an additive interaction, and scores greater than 10 reflect a synergistic drug interaction [37]. Cells were seeded in 96-well plates and treated with a concentration-combination matrix of CDDP and NFV. CDDP was administered for 3 h and then removed, after which cells were exposed to NFV for 72 h. CDDP concentrations of 2.5, 5, and 10 µM were tested in combination with NFV concentrations of 2.5, 5, 10, and 20 µM, with all concentration combinations evaluated. Cell vitality [38] was measured using the Cell Counting Kit-8 (CCK-8, Dojindo, Kumamoto, Japan) assay according to the manufacturer’s instructions, and absorbance values were recorded at 450 nm. Vitality values were normalized to vehicle-treated controls, which were set to 100% vitality. These normalized vitality data were then input to the SynergyFinder software to analyze the drug interaction.

### 2.4 Annexin-V and 7-AAD staining

Phosphatidyl serine exposure on the outer leaflet of the plasma membrane was assessed using the Muse Annexin V & Dead Cell Kit (Cytek) on the Muse Cell Analyzer according to the manufacturer’s protocol. In brief, following treatment, the floating and adherent cells were collected, centrifuged, resuspended in fresh media and stained with Annexin V & Dead Cell Reagent for 20 minutes (min) at room temperature (RT). Cell populations were quantified as live, early apoptotic, late apoptotic, and dead cells.

### 2.5 Mitochondrial membrane depolarization

Depolarization across the inner mitochondrial membrane (IMM) following drug treatment was evaluated using the Muse MitoPotential Kit (Cytek) and the Muse Cell Analyzer according to the manufacturer’s instructions. Following treatment and harvesting of floating and adherent cells, the samples were centrifuged, resuspended in 1X assay buffer, and stained with the Muse MitoPotential working solution for 20 min at 37 °C followed by incubation with the 7-AAD dye for 5 min at RT. Cells were classified by IMM polarization status and viability, and the percentage of dead/dying cells with a depolarized IMM was quantified.

### 2.6 LDH release assay

Cell death-associated plasma membrane lysis damage was assessed by measuring LDH release into culture supernatants using the Cytotoxicity Detection Kit (LDH) (Roche, Mannheim, Germany), according to the manufacturer’s instructions. Briefly, cells were seeded in 96-well plates and following treatment supernatants were collected and incubated with the LDH reaction mixture for 30 min at RT in the dark. Absorbance values were measured at 490 nm.

### 2.7 Immunoblotting

Following treatment incubation, ovarian cancer cells were collected by scraping in ice-cold 1X PBS and pelleted by centrifugation. Cell pellets were stored at -80 °C until protein extraction. Cell lysis was achieved using NP-40 lysis buffer supplemented with protease inhibitors. Lysis was done on ice for 30 min, after which lysates were clarified by centrifugation at 12,000 x *g* for 15 min at 4 °C, and protein concentrations were determined using the Pierce BCA protein assay (Thermo Fisher Scientific, Waltham, MA, USA). Twenty-five µg of total protein per lane was resolved at 200V for 35 min on 12% SDS-poly acrylamide gels prepared using the TGX Stain-Free FastCast Acrylamide Kit (Bio-Rad Laboratories, Hercules, CA, USA). Proteins were transferred onto PVDF membranes using the Trans-Blot Turbo Transfer System for 7 min (Bio-Rad). Membranes were blocked in 5% non-fat milk in TBS-T for 1 h at RT and incubated overnight at 4 °C with primary antibodies diluted 1:1000 in 5% non-fat milk in TBS-T or 5% Bovine Serum Albumin (BSA, Sigma) in TBS-T. The following primary antibodies were used for immunoblotting analysis: caspase-3 (#9622, Cell Signalling Technology, Danvers, MA, USA), PARP (#9542, Cell Signalling), gasdermin E (#84005, Cell Signalling), caspase-8 (#9746, Cell Signalling), Bid (#2002, Cell Signalling), Bcl-2 (#3498, Cell Signalling), Bax (#5023, Cell Signalling), CHOP (#2895, Cell Signalling), PUMA (#12450, Cell Signalling), phospho-AKT (Ser473) (#9271, Cell Signalling), phospho-GSK-3β (#5558, Cell Signalling), phospho-histone H2AX (Ser139) (γH2AX) (#9718, Cell Signalling), 53BP1 (#612523, BD Transduction Laboratories, Franklin Lakes, NJ, USA), and β-actin (A5441, Sigma). After washing in TBS-T, membranes were incubated with the appropriate HRP-conjugated secondary antibodies, goat anti-rabbit IgG (H+L) HRP conjugate (1706515, Bio-Rad) or goat anti-mouse IgG (H+L) HRP conjugate (1706516, Bio-Rad), for 1 h at RT. Protein bands were visualized on using Clarity Western ECL Substrate (Bio-Rad) and imaged with a ChemiDoc Touch Imaging System (Bio-Rad).

### 2.8 RNA sequencing and analysis

IGROV-1/CP cells were treated with vehicle, 10 µM CDDP for 3 h followed by culture in drug-free media for 24 h, 20 µM NFV for 24 h, or 10 µM CDDP for 3 h followed by culture in 20 µM NFV-containing media for 24 h. Cell pellets were then submitted to Genome Québec (Montreal, Quebec, Canada) for RNA extraction, library preparation, and sequencing. Total RNA was extracted using the RNeasy Plus Universal Mini Kit (QIAGEN, Hilden, Germany, cat. no. 73404) according to the manufacturer’s instructions. Briefly, the lysis buffer reagent QIAzol was added to the cell samples, and homogenization was performed by passing the lysate 15 times through a 20-gauge needle attached to a sterile 3 mL syringe. RNA was eluted in 35 µL of the buffer provided with the extraction kit. Total RNA was quantified and its integrity assessed using the 5K/RNA/Charge Variant Assay LabChip and RNA Assay Reagent Kit (PerkinElmer, Shelton, CT, USA). Libraries were generated from 250 ng of total RNA using the Illumina Stranded mRNA Prep Kit (Illumina, San Diego, CA, USA), including mRNA enrichment, according to the manufacturer’s recommendations. Libraries were quantified using the KAPA Library Quantification Kit (Kapa Biosystems, Roche), and average fragment size was determined using a Fragment Analyzer 5300 (Agilent Technologies, Santa Clara, CA, USA). Libraries were normalized, pooled, denatured in 0.02 N NaOH, neutralized with pre-load buffer, and loaded at 170 pM onto an Illumina NovaSeq X Plus 25B lane for 2 × 100 bp paired-end sequencing. A PhiX control library was spiked in at 1%. Base calling and demultiplexing were performed using BCL Convert v4.2.4 to generate FASTQ files. Gene-level count tables generated using the DRAGEN pipeline were used for downstream analysis. Differential expression analysis was performed in R using DESeq2 (v1.46.0) [39]. Principal component analysis and sample-to-sample correlation analysis were used to assess replicate clustering and consistency. Genes with an adjusted p value < 0.05 and |log2 fold change| ≥ 1 were considered significantly differentially expressed. Gene set enrichment analysis was performed in R using fgsea (v1.32.4) [40] with genes ranked by the Wald statistic from DESeq2 differential expression analysis. Hallmark and Gene Ontology Biological Process gene sets obtained from the Molecular Signatures Database (MSigDB v2025.1.Hs) [41] using msigdbr (v25.1.1). KEGG[42] pathway gene sets were retrieved using KEGGREST (v1.46.0) and analyzed using the same fgsea-based preranked enrichment workflow.

### 2.9 Statistical analysis

Data are presented as mean ± SD from three independent experiments unless otherwise indicated. Statistical analyses were performed using GraphPad Prism (Dotmatics, Boston, MA, USA). Statistical significance among groups was determined using one-way ANOVA followed by Tukey’s post hoc multiple comparisons test. Where appropriate, two-way ANOVA followed by Sidak’s multiple comparisons test was used. A *p* value, or a false discovery rate (FDR) adjusted *p* value (adj *p*), < 0.05 was considered statistically significant.

## 3. Results

### 3.1 NFV synergizes with CDDP to enhance cytotoxicity in Pt-resistant ovarian cancer cells

To determine whether NFV enhances the cytotoxic effects of CDDP in ovarian cancer cells, PEO4, IGROV-1/CP, and TOV21G cells were exposed to increasing concentrations of CDDP (0, 10, 25, 50, or 100 µM) for 3 h, followed by culture in drug-free medium or medium containing NFV (20 µM) for 72 h. Across all three cell lines, the addition of NFV enhanced the cytotoxic effects of CDDP, resulting in lower viability across the tested CDDP concentrations than with CDDP alone (Fig. 1A-C). To quantify the effect of NFV on CDDP toxicity, we compared CDDP IC50 values, defined as the concentration of CDDP required to reduce cell viability by 50%, in the absence or presence of 20 µM NFV. The addition of NFV reduced the CDDP IC50 by approximately 4-fold in PEO4 and IGROV-1/CP cells and by 5.5-fold in TOV21G cells (Fig. 1E). Synergy analysis using the Loewe additivity model [36] further supported a cooperative interaction between the two agents. Loewe synergy plots showed positive synergy across dose combinations in all three cell lines (Fig. 1D), with mean Loewe synergy scores of approximately 25-30, consistent with strong synergy in each model tested (Fig. 1F). Together, these findings show that NFV markedly enhances CDDP cytotoxicity in Pt-resistant ovarian cancer cells.

**Figure 1.**
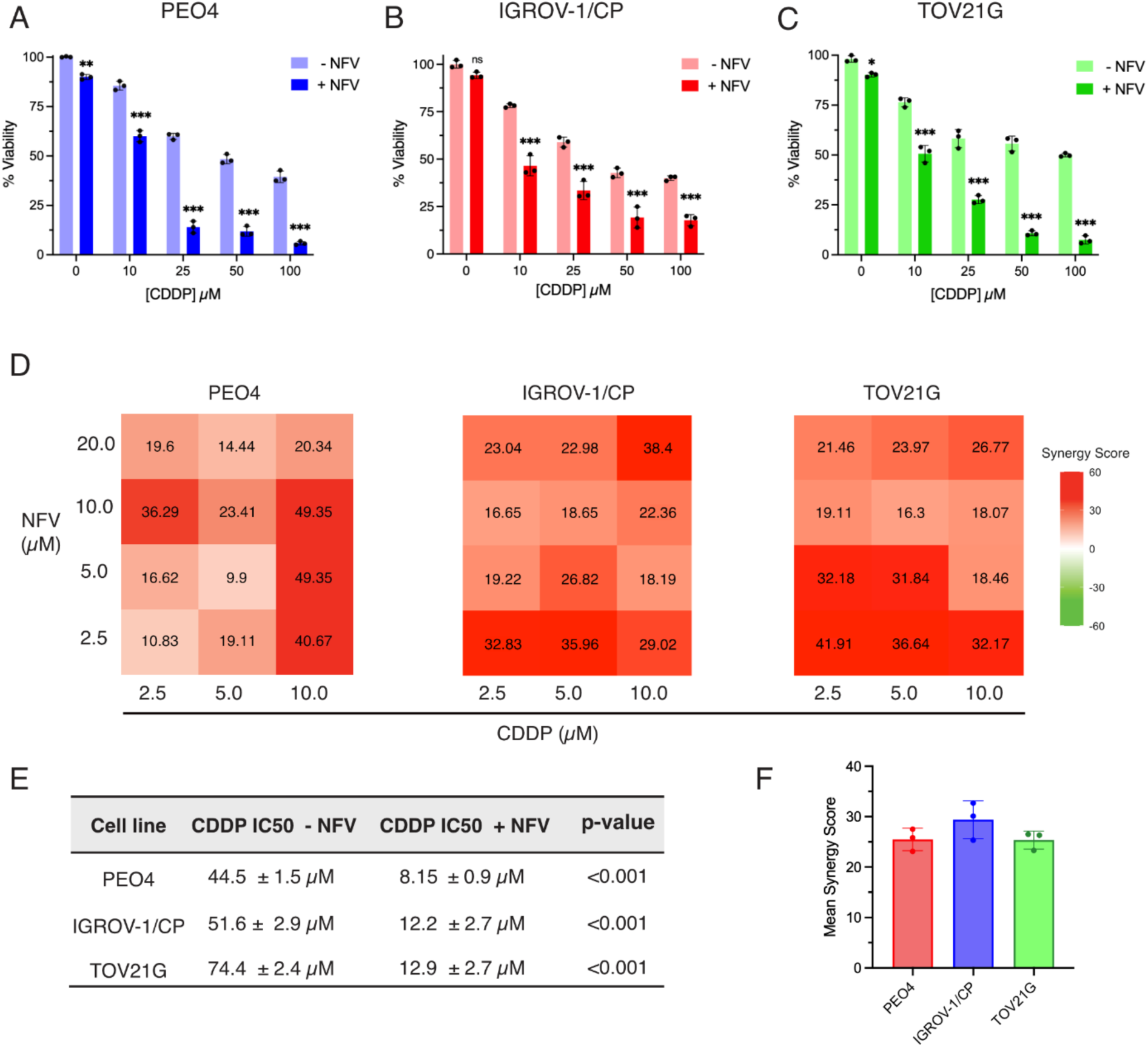
Nelfinavir synergizes with cisplatin to reduce viability in ovarian cancer cells. **(A-C)** Dose-response graphs showing cell viability following treatment with increasing concentrations of cisplatin (CDDP) alone or in combination with 20 µM nelfinavir (NFV) in PEO4, IGROV-1/CP, and TOV21G cells respectively. NFV alone is included at 0 µM CDDP. **(D)** Loewe synergy heatmaps generated using SynergyFinder analysis software, showing synergy scores for all tested CDDP/NFV dose combinations in PEO4, IGROV-1/CP and TOV21G cells. Scores less than -10 are considered to be antagonistic, scores from -10 to 10 are considered additive, and scores greater than 10 are considered synergistic. **(E)** Mean IC50 values for CDDP alone and CDDP when in combination with 20 µM NFV in each cell line. **(F)** Mean Loewe synergy scores calculated by averaging the scores from all individual dose combinations in each cell line. Data are presented as mean ± SD from three independent experiments.

### 3.2 CDDP/NFV-mediated cytotoxicity is associated with the activation of molecular mediators of apoptosis and with disruption of plasma membrane integrity, leading to cell lysis

To explore the cell death modality induced by the combination of CDDP with NFV, we assessed Annexin V/7-AAD staining, cellular morphology, LDH release, and cleavage of key cell death effectors following treatment. In both PEO4 and IGROV-1/CP cells, microcapillary-flow cytometric analysis showed that the combination treatment increased the proportion of Annexin V- and 7-AAD-positive cells relative to vehicle and single-agent controls, consistent with enhanced cell death accompanied by loss of membrane integrity (Fig. 2A,B). Representative brightfield images supported a mixed death phenotype following combination treatment; in addition to detached, shrunken cells consistent with apoptosis, PEO4 cells displayed enlarged detached cells with ballooning morphology, whereas IGROV-1/CP cells exhibited prominent vacuolization and cellular swelling, features consistent with the membrane ballooning and cell swelling commonly associated with a pyroptotic cell death phenotype [43, 44] (Fig. 2C). To further define the molecular features of this response, we examined cleavage of caspase-3 and its downstream target PARP [45], and of GSDME. In both cell lines, the drug combination increased cleaved caspase-3, cleaved PARP, and cleaved GSDME, consistent with engagement of death programs with both apoptotic and pyroptotic molecular markers (Fig. 2D,E). In parallel, the drug combination significantly increased LDH release relative to either monotherapy in both cell lines, further supporting plasma membrane rupture (Fig. 2F,G). When CDDP/NFV-treated cells were co-treated with glycine, LDH release following combination was reduced, supporting a contribution of the plasma membrane rupture to the observed death phenotype (Fig 2F,G). Glycine has long been recognized for its cytoprotective effects [46] and was recently shown to inhibit oligomerization of the plasma membrane protein NINJ1 to prevent cell lysis [47]. Collectively, these findings indicate that the combination of CDDP and NFV induces a lytic death phenotype in Pt-resistant ovarian cancer cells with activation of apoptotic and pyroptotic molecular mediators.

**Figure 2.**
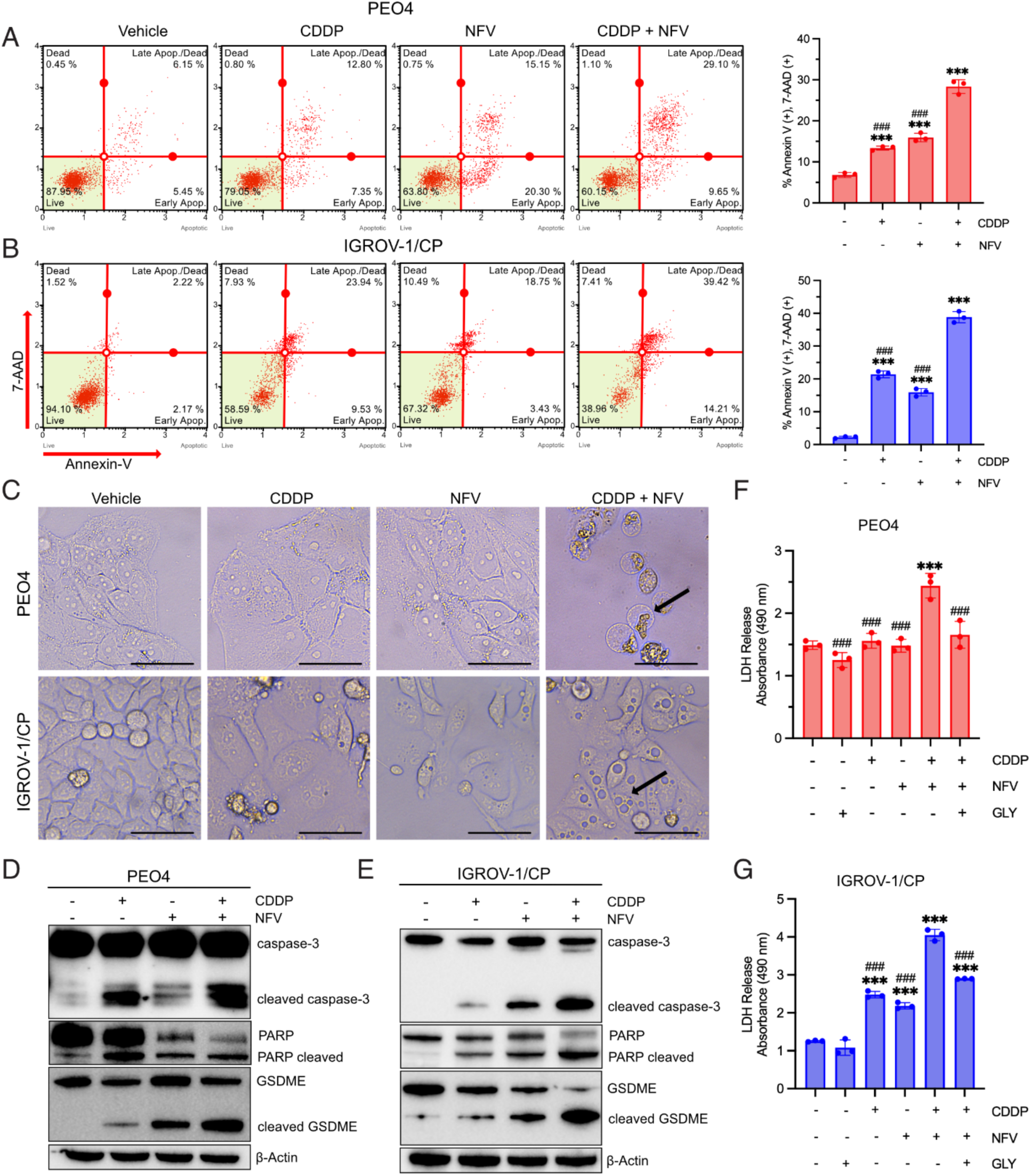
Combined nelfinavir and cisplatin treatment induces a lytic cell death phenotype with apoptotic and pyroptotic features in ovarian cancer cells. PEO4 and IGROV-1/CP cells were treated with vehicle for 72 h, cisplatin (CDDP) for 3 h followed by drug-free medium for 72 h, nelfinavir (NFV) for 72 h, or CDDP for 3 h followed by NFV for 72 h, as indicated. **(A,B)** Representative Annexin V/7-AAD microcapillary-flow cytometry plots and quantification of Annexin V-positive/7-AAD-positive cells in PEO4 (**A**) and IGROV-1/CP (**B**) cells. **(C)** Representative brightfield images of PEO4 and IGROV-1/CP cells following treatments. Black arrows indicate enlarged ballooned cells in PEO4 and swollen/vacuolated cells in IGROV-1/CP following combination treatment. Scale bars = 50 µm. **(D,E)** Immunoblot analysis of caspase-3, PARP, and gasdermin E (GSDME) cleavage in PEO4 (**D**) and IGROV-1/CP (**E**) cells following treatment. **(F,G)** Lactate dehydrogenase (LDH) release assay in PEO4 (**F**) and IGROV-1/CP (**G**) cells following CDDP/NFV treatment in the absence or presence of 5 mM glycine (GLY). Data are presented as mean ± SD from. Statistical significance was determined by one-way ANOVA with Tukey’s post hoc test. ***P < 0.001 versus vehicle; ###P < 0.001 versus CDDP plus NFV.

### 3.3 The combination treatment CDDP/NFV engages caspase-8 and mitochondrial apoptotic signalling

Given that the combination of CDDP and NFV induced cell lysis and the activation of apoptotic and pyroptotic biomarkers, we next asked whether the drugs engaged upstream apoptotic signalling pathways. Caspase-8 can couple upstream death signalling to the mitochondrial apoptotic pathway through Bid processing into cleaved Bid, a pro-apoptotic Bcl-2 family member [21]. Consistent with this, the CDDP/NFV treatment increased caspase-8 cleavage in both PEO4 and IGROV-1/CP cells relative to vehicle and single-agent controls (Fig. 3A). In parallel, we observed a reduction in full-length Bid, consistent with Bid processing downstream of caspase-8 activation (Fig 3A). To evaluate mitochondrial integrity in drug-treated cells, we examined the relative levels of the pro-apoptotic protein Bax and the anti-apoptotic protein Bcl-2, as the Bax/Bcl-2 ratio is a major determinant of outer mitochondrial membrane (OMM) permeabilization [48]. Combined, CDDP and NFV increased the Bax/Bcl-2 ratio in both cell lines, indicating a shift toward a pro-apoptotic mitochondrial state (Fig. 3B) [48]. In agreement with this conclusion, analysis of the IMM potential revealed an increased proportion of cells with a depolarized IMM following combination treatment (Fig. 3C,D). Transcriptomic analysis further supported activation of these apoptotic programs. Gene ontology biological process enrichment [49, 50] demonstrated significant upregulation of extrinsic apoptotic signalling, intrinsic apoptotic signalling, and apoptotic mitochondrial changes in IGROV-1/CP cells treated with both CDDP and NFV relative to vehicle-treated controls (Fig. 3E). Together, these findings indicate that, when combined, CDDP and NFV engage caspase-8-associated mitochondrial apoptotic signalling in Pt-resistant ovarian cancer cells.

**Figure 3.**
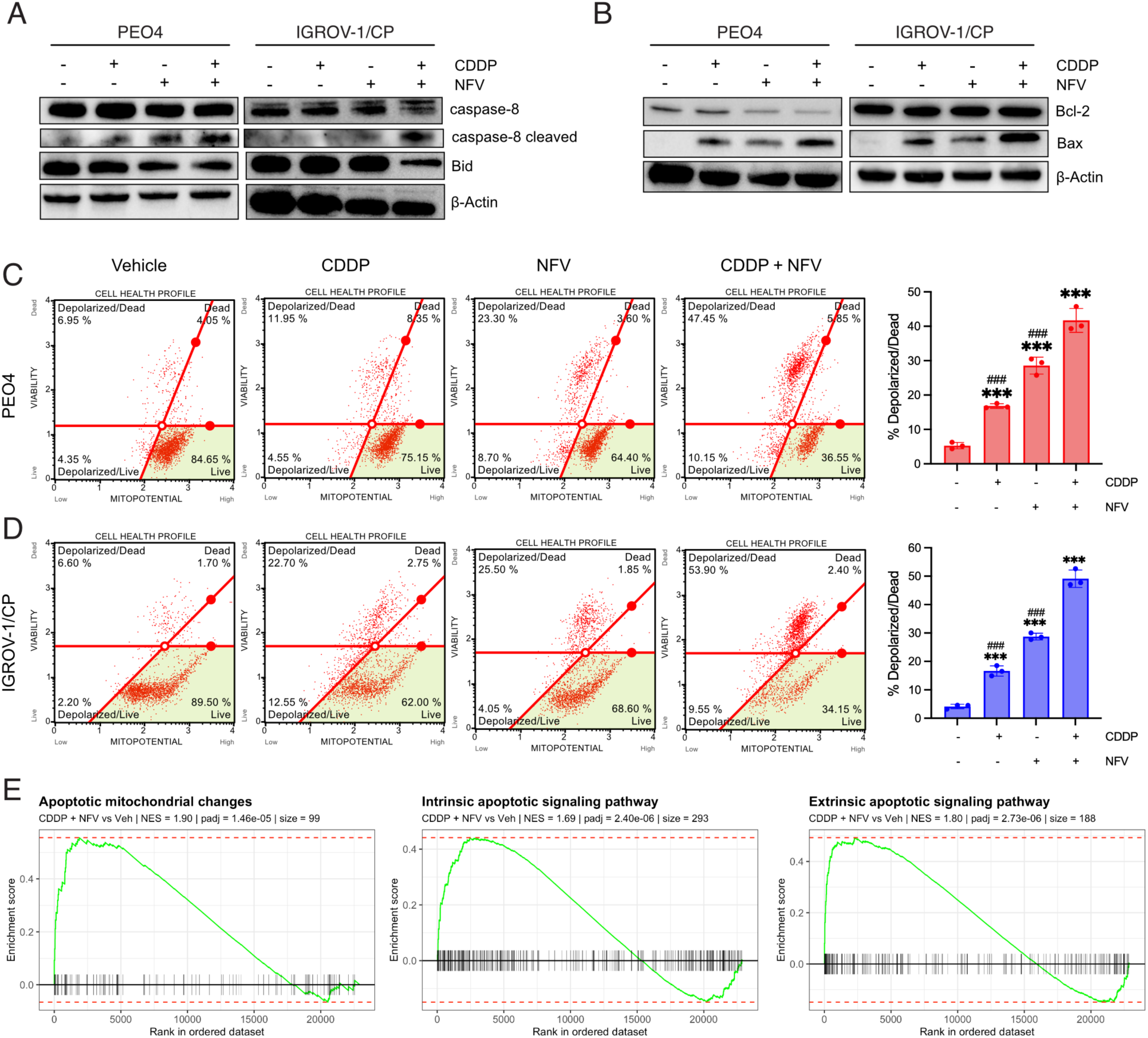
Combined nelfinavir and cisplatin treatment engages caspase-8-associated mitochondrial apoptotic signalling in ovarian cancer cells. PEO4 and IGROV-1/CP cells were treated with vehicle (72 h), CDDP (3 h) followed by drug-free medium (72 h), NFV (72 h), or CDDP (3 h) followed by NFV (72 h), as indicated. **(A)** Immunoblot analysis of caspase-8 and Bid in PEO4 and IGROV-1/CP cells following treatment. **(B)** Immunoblot analysis of Bcl-2 and Bax in PEO4 and IGROV-1/CP cells following treatment. β-Actin was used as a loading control. **(C,D)** Representative mitochondrial membrane potential microcapillary-flow cytometry plots and quantification of depolarized/dead cells in PEO4 (**C**) and IGROV-1/CP (**D**) cells following treatment. **(E)** Gene set enrichment plots showing enrichment of extrinsic apoptotic signalling, intrinsic apoptotic signalling, and apoptotic mitochondrial changes in CDDP/NFV-treated IGROV-1/CP cells relative to vehicle controls. Data are presented as mean ± SD. Statistical significance was determined by one-way ANOVA with Tukey’s post hoc test. ***P < 0.001 versus vehicle; ###P < 0.001 versus CDDP/NFV.

### 3.4 CDDP/NFV-induced cell death depends on caspase-8 and caspase-3, but does not involve necroptosis or ferroptosis

Having identified activation of caspase-8-associated mitochondrial apoptotic signalling together with caspase-3 and GSDME cleavage in response to CDDP/NFV, we next examined whether these caspases were functionally required for the regulated lytic cell death observed. In both PEO4 and IGROV-1/CP cells, inhibition of caspase-3 with Z-DEVD-fmk significantly rescued viability following the combination treatment, indicating that caspase-3 activation is required to drive cell death (Fig. 4A). Similarly, inhibition of caspase-8 with Z-IETD-fmk upon treatment with the CDDP and NFV combination improved viability in both cell lines, supporting a functional role for caspase-8 upstream of this response (Fig. 4B). Immunoblot analysis further supported this signalling hierarchy. Inhibition of caspase-3 with Z-DEVD-fmk prevented GSDME cleavage following CDDP/NFV treatment, indicating that GSDME processing occurs downstream of caspase-3 activation (Fig. 4C). In addition, inhibition of caspase-8 with Z-IETD-fmk reduced cleavage of both caspase-3 and GSDME, placing caspase-8 upstream of the apoptotic and lytic machinery engaged by the drug combination (Fig. 4D). Because lytic cell death can result from regulated cell death pathways other than pyroptosis, we next asked whether ferroptosis and necroptosis contributed to the cell lysis observed following CDDP/NFV treatment. To test this, cells were treated with the combination of CDDP and NFV in the presence of ferrostatin-1, a radical-trapping antioxidant that inhibits lipid peroxidation to prevent ferroptosis [51], or necrostatin-1, an inhibitor of RIPK1-dependent necroptosis [52]. Neither inhibitor rescued cell viability in either cell line following CDDP/NFV treatment when used at concentrations reported to block ferroptosis and necroptosis [51, 52] in cancer cells (Fig. 4E,F). Collectively, these findings demonstrate that the cell lysis induced by CDDP/NFV depends on activation of both caspase-8 and caspase-3, is associated with GSDME cleavage, and does not involve activation of either ferroptosis or necroptosis.

**Figure 4.**
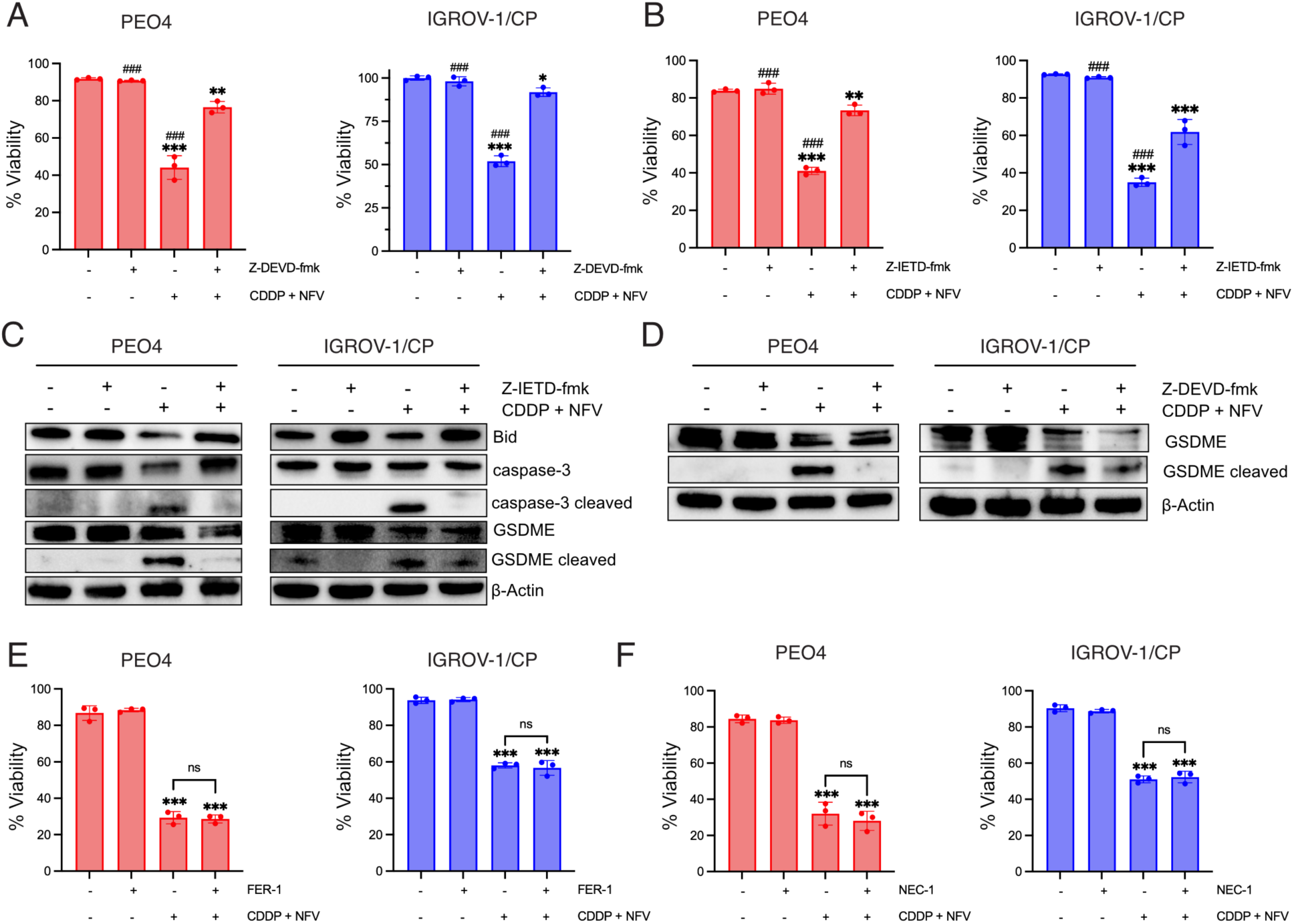
CDDP/NFV-induced cell death depends on caspase-8 and caspase-3 and does not involve ferroptosis or necroptosis. PEO4 and IGROV-1/CP cells were treated with vehicle or CDDP (10 µM, 3 h) followed by NFV (20 µM, 72 h), in the absence or presence of Z-DEVD-fmk (caspase-3 inhibitor; 100 µM), Z-IETD-fmk (caspase-8 inhibitor; 100 µM), ferrostatin-1 (FER-1, ferroptosis inhibitor; 1 µM), or necrostatin-1 (NEC-1, necroptosis inhibitor; 10 µM), as indicated. **(A,C)** Cell viability in PEO4 and IGROV-1/CP cells following treatment with CDDP + NFV in the absence or presence of Z-DEVD-fmk (**A**) or Z-IETD-fmk (**C**). **(B)** Immunoblot analysis of Bid, caspase-3, and gasdermin E (GSDME) in PEO4 and IGROV-1/CP cells treated with CDDP + NFV in the absence or presence of Z-IETD-fmk. **(D)** Immunoblot analysis of GSDME cleavage in PEO4 and IGROV-1/CP cells treated with CDDP/ NFV in the absence or presence of Z-DEVD-fmk. **(E,F)** Cell viability in PEO4 and IGROV-1/CP cells following treatment with CDDP/ NFV in the absence or presence of FER-1 (**E**) or NEC-1 (**F**). Data are presented as mean ± SD. Statistical significance was determined by one-way ANOVA with Tukey’s post hoc test. *P < 0.05, **P < 0.01, and ***P < 0.001 versus vehicle; ###P < 0.001 versus CDDP /NFV; ns, not significant.

### 3.5 The combination of CDDP and NFV induces ER stress and DNA damage-associated pro-apoptotic mediators while supressing the Akt survival pathway and the DNA repair factor 53BP1

To place the apoptosis-focused transcriptomic analysis shown in Figure 3 in a broader transcriptional context, we next examined differential gene expression and Hallmark pathway enrichment in IGROV-1/CP cells treated with CDDDP/NFV relative to vehicle controls. Principal component analysis and sample-to-sample correlation analysis demonstrated appropriate replicate clustering and overall consistency across samples (Supplementary Fig. S1). At the pathway level, Hallmark analysis revealed enrichment of stress- and death-associated pathways, including reactive oxygen species (ROS), apoptosis, the unfolded protein response (UPR), and TNF/NF-κB signalling (Fig. 5A). Interestingly, the p53 signalling pathway was also enriched despite the mutant *TP53* background of IGROV-1/CP cells (Fig 5A). In contrast, pathways linked to glycolysis, cholesterol homeostasis, fatty acid metabolism, mitotic spindle, G2/M checkpoint, and E2F targets were negatively enriched (Fig. 5A). Consistent with this broader stress-associated response, ROS production was observed following combination treatment in both PEO4 and IGROV-1/CP cells (Supplementary Fig. S2). Overall, these data suggest that the combination CDDP/NFV shifts ovarian cancer cells away from metabolic and proliferative programs and toward a coordinated stress-to-death transcriptional state. To examine these responses at the gene level, we generated an integrated heatmap of selected transcripts across vehicle-, CDDP-, NFV-, and CDDP/NFV-treated groups. Genes associated with endoplasmic reticulum (ER) stress, including *DDIT3* [53] (coding for CHOP), *ATF3* [54], and *XBP1* [55], were elevated following combination treatment (Fig. 5B). Likewise, genes linked to DNA damage-associated pro-apoptotic signalling, including *GADD45A/B* [56], *BBC3* (coding for the pro-apoptotic protein PUMA) [57], and *AEN* [58], were also upregulated (Fig. 5B). Notably, we also observed increased expression of transcripts related to extrinsic apoptotic and death receptor signalling, including *TNFRSF10A* (coding for Death Receptor 4; DR4) [59], *TNFRSF10B* (coding for Death Receptor 5; DR5) [59], *TNFRSF21* (coding for Death Receptor 6; DR6) [60], and *FADD* [59] (Fig. 5B). These transcriptional changes provide further support for the activation of a caspase-8-associated death program observed at the protein and functional levels. Given that NFV has previously been linked to the suppression of pro-survival signalling, particularly through the inhibition of the PI3K-Akt pathway [15], we next examined whether these transcriptomic data support downregulation of cell survival and DNA repair pathways in our model. KEGG gene set enrichment analysis demonstrated significant negative enrichment of the PI3K-Akt signalling pathway, and of the homologous recombination (HR) and Fanconi anemia DNA repair pathways in the CDDP/NFV combination-treated cells relative to vehicle controls (Fig. 5C). These findings suggest that CDDP/NFV not only promote stress- and death-associated signalling but may also diminish adaptive survival signalling and impair pathways involved in repair of Pt-induced DNA damage. Protein-level analysis further reinforced the transcriptomic findings. Immunoblotting confirmed that CDDP/NFV treatment increased the expression of the ER-stress induced protein CHOP and its downstream transcriptional target PUMA [61] relative to vehicle and single-agent controls in both PEO4 and IGROV-1/CP cells (Fig. 5D). In parallel, the drug combination reduced Ser473 phosphorylation of Akt [62] and of its downstream target GSK-3β [62], consistent with suppression of PI3K-Akt survival signalling (Fig. 5E). Finally, we also observed a decrease in the protein levels of 53BP1, a master regulator of DNA double-strand break repair that promotes non-homologous end joining (NHEJ) following DNA damage [63], by NFV and the combination treatment. In line with this, phosphorylation of histone 2AX (γH2AX), a marker of double-strand DNA damage [64], was increased in the combination compared to vehicle and single agent controls, supporting the possibility that NFV may enhance CDDP-mediated DNA damage accumulation by limiting the ability of cells to mount an effective DNA repair response (Fig. 5F). Together, these transcriptomic and protein-level changes support a model in which the combination CDDP/NFV establishes a cellular state characterized by ER stress, increased DNA damage with diminished DNA repair, apoptotic priming, and death receptor-associated signalling, while impairing pro-survival programs.

**Figure 5.**
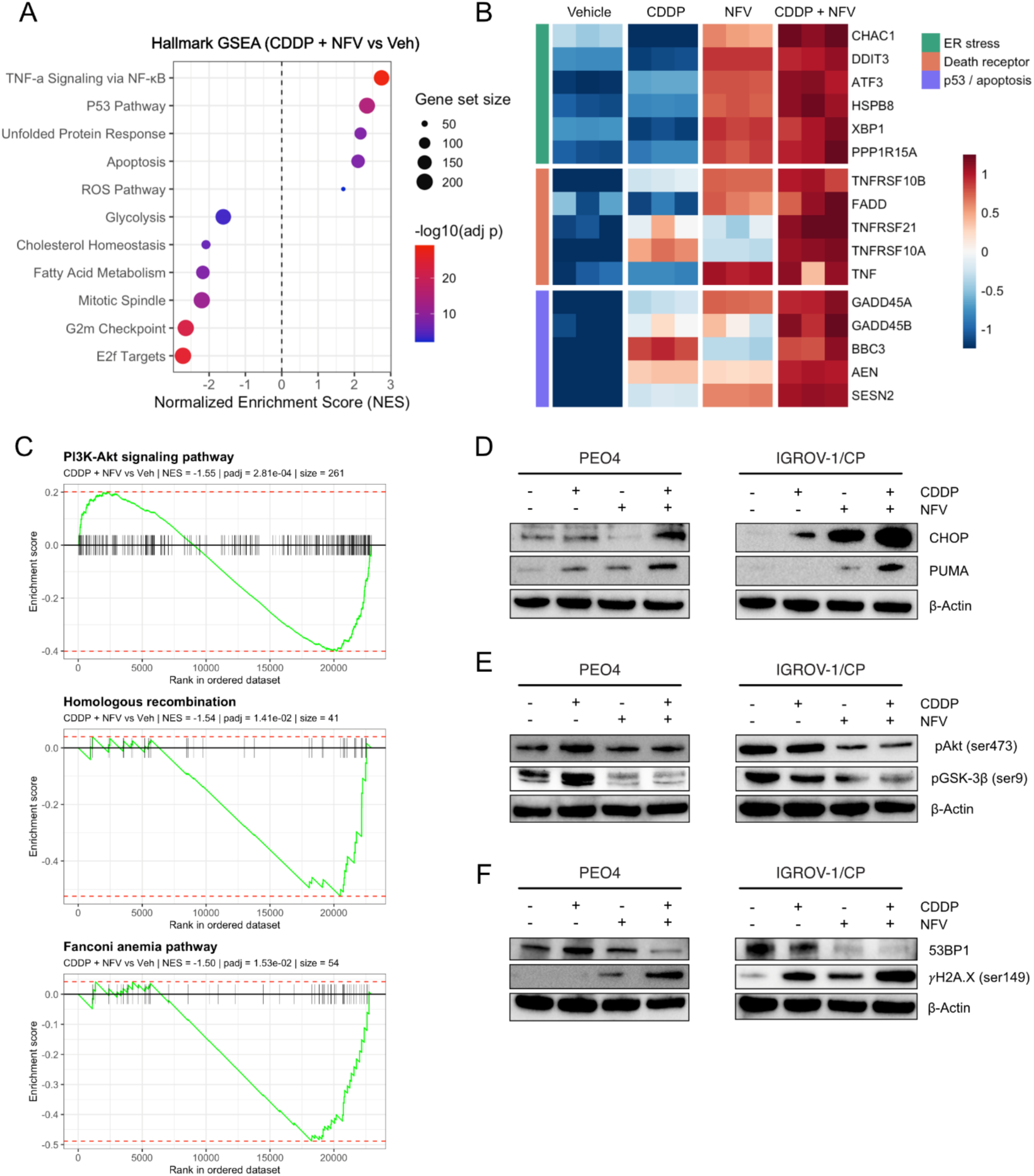
Combined nelfinavir and cisplatin treatment induces ER stress, DNA damage-associated pro-apoptotic signalling, suppression of survival and DNA repair pathways, and extrinsic apoptotic signalling programs in ovarian cancer cells. PEO4 and IGROV-1/CP cells were treated with vehicle (72 h), cisplatin (CDDP; 10 µM, 3 h) followed by drug-free medium (72 h), nelfinavir (NFV;20 µM, 72 h), or CDDP (10 µM, 3 h) followed by NFV (20 µM, 72 h), as indicated. **(A)** Hallmark gene set enrichment analysis of CDDP/NFV-treated IGROV-1/CP cells relative to vehicle controls. Dot size represents gene set size, and colour represents the -log10 adjusted p value. **(B)** Heatmap showing Z-score-scaled expression of selected transcripts associated with ER stress, death receptor signalling, and p53/apoptosis across treatment groups in IGROV-1/CP cells. Columns represent individual biological replicates. Expression values are shown as row-scaled Z scores, with the colour scale capped at ± 1.25 **(C)** Gene set enrichment plots of selected KEGG pathways in CDDP plus NFV-treated IGROV-1/CP cells relative to vehicle controls. Normalized enrichment scores (NES) adjusted p-value, and gene set size are indicated. **(D)** Representative immunoblot analysis of CHOP and PUMA expression in PEO4 and IGROV-1/CP cells following treatment. **(E)** Representative immunoblot analysis of phosphorylated Akt (Ser473) and phosphorylated GSK-3β (ser9) in PEO4 and IGROV-1/CP cells following treatment. **(F)** Representative immunoblot analysis of 53BP1 and γH2AX (ser149) in PEO4 and IGROV-1/CP cells following treatment. β-actin was used as a loading control for the immunoblot analyses.

## 4. Discussion

In this study, we show that the HIV protease inhibitor NFV markedly enhances CDDP-induced cytotoxicity in Pt-resistant ovarian cancer cells and shifts the resulting death response toward a caspase-8/caspase-3/GSDME-linked lytic phenotype. Across three Pt-resistant ovarian cancer cell lines, NFV reduced the CDDP IC50 and shows strong synergy with CDDP. Mechanistically, the CDDP/NFV combination induces features of both apoptotic and pyroptotic cell death, including Annexin V/7-AAD positivity, caspase-3 and PARP cleavage, GSDME cleavage, LDH release, and plasma membrane-disruptive morphology. We further identify caspase-8 as an upstream contributor to this response, acting before Bid loss, caspase-3 activation, and GSDME processing, in a cellular context marked by ER stress, DNA damage-associated pro-apoptotic signalling, death receptor-associated transcriptional changes, impaired DNA repair signalling, and reduced pro-survival pathway activity. Together, these findings support a model in which NFV enhances CDDP toxicity by amplifying treatment-associated cellular stress while promoting a lytic cell death response (Fig. 6).

**Figure 6.**
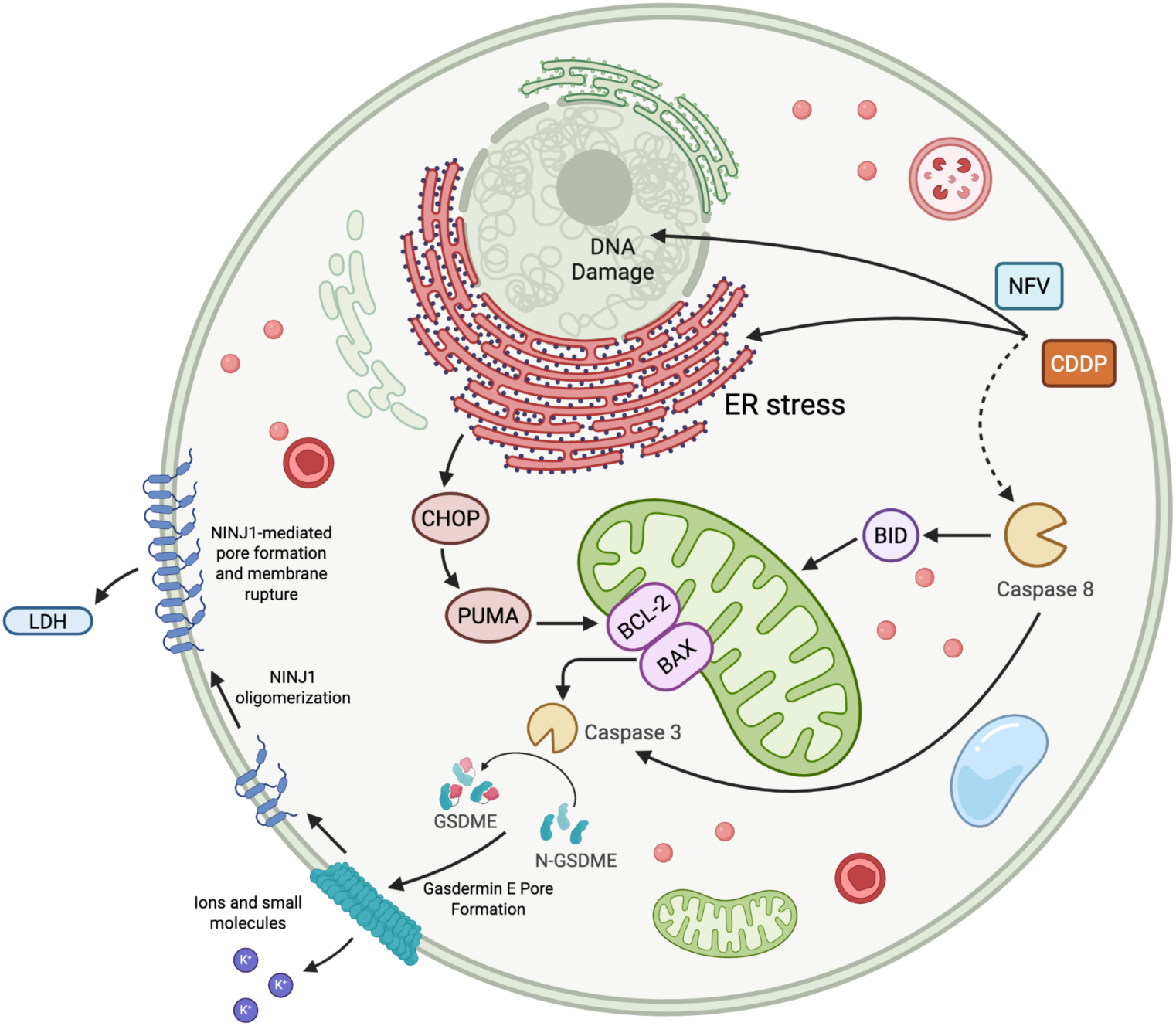
Schematic model of CDDP/NFV-induced caspase-dependent lytic cell death in Pt-resistant ovarian cancer cells. CDDP promotes DNA damage, while NFV contributes to unresolved ER stress, decreased DNA-repair capacity, and decreased cell survival signalling. Together, these cellular stress responses induce CHOP and the pro-apoptotic protein PUMA, promote a shift in the Bax/Bcl-2 balance, driving mitochondrial dysfunction. In parallel, the combination treatment engages caspase-8-associated signalling, including Bid processing, further linking upstream apoptotic signalling to the mitochondrial apoptotic pathway. These events converge on activation of caspase-3, which cleaves GSDME to generate N-terminal GSDME, leading to pore formation in the plasma membrane and the release of ions and other small intracellular molecules. This in turn is followed by oligomerization of NINJ1, triggering membrane rupture and allowing for the release of LDH culminating in complete cell lysis.

These findings are particularly relevant in ovarian cancer, where Pt-based chemotherapy remains a cornerstone of treatment but is frequently undermined by recurrence and the eventual emergence of Pt-resistant disease [11]. In many cases, Pt-resistance reflects not simply a failure to induce damage, but also an impaired ability to convert cellular stress and damage signals into the efficient execution of cell death programs [4, 65]. In this context, drug repurposing strategies that intensify responses to Pt-derivatives with clinically available agents are especially attractive [66]. NFV has emerged as a promising repurposing candidate because, beyond its original antiviral role, it has been linked to ER stress induction, disruption of proteostatic homeostasis, inhibition of pro-survival pathways, and enhanced responsiveness to anticancer therapy in multiple model systems, including high-grade serous ovarian cancer [14, 15, 67, 68]. Our findings extend this rationale in Pt-resistant ovarian cancer cells by showing that NFV not only enhances the cytotoxicity of CDDP in a synergistic manner but also alters the nature of the death response that follows. This could be especially relevant in tumors that sustain Pt-induced toxicity yet fail to convert such toxicity into an efficient cell death program.

Our data suggest that NFV helps to lower the threshold for cell death commitment by reinforcing stress pathways that complement CDDP-induced genotoxic injury, while weakening pathways that would otherwise permit survival and recovery. CDDP is a potent DNA-damaging agent [9, 18], yet many ovarian cancer cells survive CDDP exposure by mounting adaptive responses that blunt apoptotic execution [4, 69]. Our data suggest that NFV adds a complementary layer of cellular stress that helps tip this balance toward cell death execution involving not only apoptotic but also pyroptotic cell death molecular mediators. Transcriptomic and protein-level analyses showed activation of the UPR, increased expression of ER stress markers such as CHOP and GRP78, enhanced DNA damage-associated signalling, as indicated by elevated γH2AX, and induction of pro-apoptotic mediators such as PUMA. Pathway enrichment further supported increased apoptotic signalling, ROS-related signalling, and p53-associated transcriptional output, together with suppression of proliferative, metabolic, DNA repair, and PI3K-Akt associated survival programs. Collectively, these findings support a model in which the addition of NFV to CDDP shifts cells from a sublethal or incompletely executed apoptotic response into a terminal caspase-8/-3/GSDME-driven secondary pyroptotic death program. This is notable, as GSDME has increasingly emerged as a determinant of how chemotherapy-induced caspase-3 activation is ultimately expressed: in cells with sufficient GSDME, executioner caspases can drive membrane pore formation and terminal lysis rather than a non-lytic apoptotic endpoint [70]. Cells that do not complete apoptosis may persist within the tumor with substantial damage and cellular stress, whereas channeling caspase-3 activity into GSDME-associated secondary pyroptosis may make the tumor cells less recoverable [70–72]. Related work in lung adenocarcinoma has also linked NFV-based treatment to caspase-3 and GSDME cleavage, suggesting that engagement of this axis may extend beyond ovarian cancer, although the upstream context may differ across tumor types [73].

One of the most important findings in this study is the upstream role of caspase-8. Combination CDDP/NFV treatment enhanced caspase-8 cleavage whereas pharmacologic inhibition of caspase-8 improved viability while reducing caspase-3 and GSDME cleavage, placing caspase-8 upstream of both the apoptotic/pyroptotic execution and the lytic branch of the response. This is of particular interest given that caspase-8 sits at the interface of extrinsic and intrinsic death signalling pathways. Through Bid processing, caspase-8 can amplify mitochondrial apoptosis while also contributing directly to executioner caspase activation [74]. The reduction in full-length Bid, the increase in the Bax/Bcl-2 ratio, and the mitochondrial depolarization we observed here are consistent with this type of amplification [48]. In parallel, RNA-seq analysis of CDDP/NFV-treated Pt-resistant ovarian cancer cells revealed increased expression of death receptor-associated transcripts, including DR4, -5, and -6, and FADD, raising the possibility that the CDDP/NFV combination treatment establishes a cellular state permissive for death receptor-linked or death receptor-like caspase-8 activation [75, 76]. CHOP may be particularly relevant in this context, as unresolved ER stress can increase DR5 expression through CHOP activity and promote death-inducing signalling complex (DISC) formation, thereby linking the ER stress response to the extrinsic apoptotic signalling pathway [77, 78]. However, the precise upstream trigger remains unresolved. ER stress, ROS accumulation, death receptor priming, and ligand-independent death receptor signalling may each contribute to the activation of caspase-8 and warrant further investigation [76, 79–81]. Even so, the identification of caspase-8 as a functional upstream node provides an important mechanistic framework for understanding how CDDP and NFV cooperate to intensify downstream caspase-3/GSDME signalling.

Another interesting aspect of the response of the Pt-resistant ovarian cancer cells to the combination CDDP/NFV was the emergence of a p53-associated pro-apoptotic transcriptional program despite the cells’ mutant p53 background. This is particularly relevant in ovarian cancer, where *TP53* mutations and loss of p53 function are defining features in the majority of cases [1, 82, 83]. The drug combination treatment enriched the p53 pathway, as evidenced by increased expression of transcripts such as *BBC3*, *GADD45A/B*, and *AEN* all of which are commonly associated with canonical p53-mediated cellular stress responses [84]. Rather than indicating restoration of wild-type p53 function, these findings more likely reflect activation of stress-responsive transcriptional circuitry that converges on components of the classical p53 output. CHOP may again be relevant in this context. In addition to serving as a marker of unresolved ER stress, CHOP can promote apoptotic priming through induction of BH3-only proteins, such as PUMA, BIM, and NOXA [53, 61, 81, 85]. Consistent with this idea, increased PUMA expression was also observed at the protein level across both p53-mutant cell lines examined. Thus, the addition of NFV to CDDP treatment may help ovarian cancer cells acquire a ‘p53-like’ pro-apoptotic state through alternative stress pathways, potentially bypassing an important barrier to effective CDDP-induced death in *TP53*-mutant ovarian cancer cells.

Finally, our data also suggest that NFV contributes to the drug combination response by suppressing adaptive survival signalling and impairing DNA damage repair capacity. At the protein level, NFV reduced phosphorylation of Akt and its downstream target GSK-3β, with transcriptomic analysis confirming negative enrichment of PI3K-Akt signalling by the combination CDDP/NFV. This is likely important as Akt activity has been linked to CDDP resistance by promoting cell survival, sustaining metabolic adaptation, and opposing apoptotic execution [10, 86]. Our data are supported by other studies done in ovarian cancer, in which activation of the PI3K-Akt pathway has been associated with reduced Pt-sensitivity and persistence of damaged cells despite ongoing genotoxic stress, while inhibition of Akt was shown to resensitize resistant cells to CDDP [87, 88]. In parallel, CDDP/NFV treatment increased γH2AX levels, while NFV, alone and in combination with CDDP, reduced 53BP1 expression, consistent with an accumulation of unresolved DNA damage in the setting of hindered DNA repair capacity [89]. In addition to loss of 53BP1 loss, we observed negative enrichment of HR and Fanconi anemia pathways, both of which are highly important for the repair of CDDP-induced DNA lesions [90, 91]. This is particularly important in ovarian cancer, where restoration of HR capacity, including through BRCA reversion mutations that restore wild-type function such as those reported in PEO4 cells [92], is a recognized mechanism of acquired Pt-resistance [93]. In this context, NFV-mediated suppression of repair-associated factors, such as 53BP1, together with downregulation of HR and Fanconi anemia signalling, may help limit the ability of Pt-resistant cells to recover from CDDP-induced DNA damage. Together, these effects provide a plausible explanation for how NFV helps convert Pt-induced stress into a less recoverable death response.

Several limitations of this study should be acknowledged. First, these findings are based on in vitro Pt-resistant ovarian cancer models, and it remains to be determined whether NFV promotes similar death programs in vivo, particularly within the context of the tumor microenvironment. Second, although our data identify caspase-8 as a functional upstream mediator, they do not establish the precise molecular event responsible for its activation. Finally, although our pharmacologic experiments did not support major contributions from necroptosis or ferroptosis under the conditions tested, context-dependent involvement of other regulated death pathways cannot be fully excluded. This may be particularly relevant for ferroptosis, as recent work has shown that NFV alone can induce ER stress-associated ferroptotic cell death in liver cancer cells [94], suggesting that the dominant death program engaged by NFV may vary according to the cellular background and whether NFV is used as a single agent or in combination with chemotherapy [94]. Despite these limitations, our results provide a mechanistic basis for considering NFV as a Pt-enhancing partner to further cytotoxicity, especially in cases of Pt-resistant ovarian cancers. By amplifying cellular stress signalling and promoting caspase-8/caspase-3/GSDME-linked lytic cell death, NFV may enhance CDDP efficacy in tumors that would otherwise evade complete death execution, supporting its further evaluation as a repurposed therapeutic adjunct in Pt-treated ovarian cancers.

## Ethics Declaration

Not applicable.

## CRediT Author Statement

**Benjamin Forgie:** Conceptualization, Writing-original draft, Methodology, Investigation, Formal analysis, Visualization. **Rewati Prakash:** Investigation, Writing-Review and Editing. **Desiree Marno:** Investigation, Writing-Review and Editing. **Farah H. Abdalbari:** Investigation, Writing-Review and Editing. **Edith Zorychta:** Writing – Review and Editing. **Abu Shadat M. Noman:** Funding acquisition. **Alicia A. Goyeneche:** Writing-Review and Editing, Supervision, Methodology. **Lucy Gilbert:** Funding acquisition, Resources, Writing-Review and Editing. **Julia V. Burnier:** Writing-Review and Editing, Supervision, Funding acquisition, Resources. **Carlos M. Telleria:** Writing-Review and Editing, Supervision, Resources, Project Administration, Funding Acquisition, Conceptualization.

## Acknowledgements

This work was supported by a grant from DxQuest Inc. North America, Ovarian Cancer Canada, internal funds from the Gerald Bronfman Department of Oncology at McGill University (to LG) and a Canada Graduate Scholarship Doctoral Research Award (CGSD) awarded to Benjamin Forgie (award #193306). JVB was supported by Fonds de Recherche du Québec en Santé (award #312831).

## Data availability

Data will be made available on request.

## References

1 M.A. Lisio, L. Fu, A. Goyeneche, Z.H. Gao and C. Telleria, Int J Mol Sci 20, (2019) doi: 10.3390/ijms20040952

2 B.M. Reid, J.B. Permuth and T.A. Sellers, Cancer Biol Med 14, 9–32 (2017) doi: 10.20892/j.issn.2095-3941.2016.0084

3 L. Wang, Q. Zhang, X. Wang, Z. Dong, S. Liu, Q. Wang, Z. Zhang and J. Xing, Biomark Res 13, 103 (2025) doi: 10.1186/s40364-025-00818-7

4 H. Li, J.J. Sheng, S.A. Zheng, P.W. Liu, N. Wu, W.J. Zeng, Y.H. Li and J. Wang, Genes Dis 13, 101801 (2026) doi: 10.1016/j.gendis.2025.101801

5 C.P. Huang, M. Fofana, J. Chan, C.J. Chang and S.B. Howell, Metallomics 6, 654–661 (2014) doi: 10.1039/c3mt00331k

6 H. Masuda, R.F. Ozols, G.-M. Lai, A. Fojo, M. Rothenberg and T.C. Hamilton, Cancer Research 48, 5713–5716 (1988)

7 D. Criscuolo, R. Avolio, M. Parri, S. Romano, P. Chiarugi, D.S. Matassa and F. Esposito, Antioxidants (Basel) 11, (2022) doi: 10.3390/antiox11081544

8 L.J. Bao, M.C. Jaramillo, Z.B. Zhang, Y.X. Zheng, M. Yao, D.D. Zhang and X.F. Yi, Int J Clin Exp Pathol 7, 1502–1513 (2014)

9 B.N. Forgie, R. Prakash and C.M. Telleria, Int J Mol Sci 23, (2022) doi: 10.3390/ijms232315410

10 Z.N. Navaei, G. Khalili-Tanha, A.S. Zangouei, M.R. Abbaszadegan and M. Moghbeli, Oncol Res 29, 235–250 (2021) doi: 10.32604/or.2022.025323

11 A. Zoń and I. Bednarek, Int J Mol Sci 24, (2023) doi: 10.3390/ijms24087585

12 C.M. Perry and P. Benfield, Drugs 54, 81–87 (1997) doi: 10.2165/00003495-199754010-00007

13 V.S. Kulkarni, V. Alagarsamy, V.R. Solomon, P.A. Jose and S. Murugesan, Russ J Bioorg Chem 49, 157–166 (2023) doi: 10.1134/s1068162023020139

14 M.R. Subeha and C.M. Telleria, Cancers 12, 3437 (2020)

15 M.R. Subeha, A.A. Goyeneche, P. Bustamante, M.A. Lisio, J.V. Burnier and C.M. Telleria, Cancers (Basel) 14, (2021) doi: 10.3390/cancers14010099

16 A. Fundytus, M. Sengar, D. Lombe, W. Hopman, M. Jalink, B. Gyawali, D. Trapani, F. Roitberg, E.G.E. De Vries, L. Moja, A. Ilbawi, R. Sullivan and C.M. Booth, The Lancet Oncology 22, 1367–1377 (2021) doi: 10.1016/S1470-2045(21)00463-0

17. A. Mariconda, J. Ceramella, A. Catalano, C. Saturnino, M.S. Sinicropi and P. Longo, in Inorganics, (2025), p. 246

18 S. Dasari and P.B. Tchounwou, Eur J Pharmacol 740, 364–378 (2014) doi: 10.1016/j.ejphar.2014.07.025

19 A. Basu and S. Krishnamurthy, J Nucleic Acids 2010, (2010) doi: 10.4061/2010/201367

20 I. Budihardjo, H. Oliver, M. Lutter, X. Luo and X. Wang, Annu Rev Cell Dev Biol 15, 269–290 (1999) doi: 10.1146/annurev.cellbio.15.1.269

21 H. Li, H. Zhu, C.J. Xu and J. Yuan, Cell 94, 491–501 (1998) doi: 10.1016/s0092-8674(00)81590-1

22 M. Fritsch, S.D. Günther, R. Schwarzer, M.-C. Albert, F. Schorn, J.P. Werthenbach, L.M. Schiffmann, N. Stair, H. Stocks, J.M. Seeger, M. Lamkanfi, M. Krönke, M. Pasparakis and H. Kashkar, Nature 575, 683–687 (2019) doi: 10.1038/s41586-019-1770-6

23 M. Jiang, L. Qi, L. Li and Y. Li, Cell Death Discovery 6, 112 (2020) doi: 10.1038/s41420-020-00349-0

24 Y. Wang, W. Gao, X. Shi, J. Ding, W. Liu, H. He, K. Wang and F. Shao, Nature 547, 99–103 (2017) doi: 10.1038/nature22393

25 J. Ding, K. Wang, W. Liu, Y. She, Q. Sun, J. Shi, H. Sun, D.-C. Wang and F. Shao, Nature 535, 111–116 (2016) doi: 10.1038/nature18590

26 P. Broz, Seminars in Immunology 69, 101811 (2023) doi: 10.1016/j.smim.2023.101811

27 N. Kayagaki, O.S. Kornfeld, B.L. Lee, I.B. Stowe, K. O’Rourke, Q. Li, W. Sandoval, D. Yan, J. Kang, M. Xu, J. Zhang, W.P. Lee, B.S. McKenzie, G. Ulas, J. Payandeh, M. Roose-Girma, Z. Modrusan, R. Reja, M. Sagolla, J.D. Webster, V. Cho, T.D. Andrews, L.X. Morris, L.A. Miosge, C.C. Goodnow, E.M. Bertram and V.M. Dixit, Nature 591, 131–136 (2021) doi: 10.1038/s41586-021-03218-7

28 M. Degen, J.C. Santos, K. Pluhackova, G. Cebrero, S. Ramos, G. Jankevicius, E. Hartenian, U. Guillerm, S.A. Mari, B. Kohl, D.J. Müller, P. Schanda, T. Maier, C. Perez, C. Sieben, P. Broz and S. Hiller, Nature 618, 1065–1071 (2023) doi: 10.1038/s41586-023-05991-z

29 Y. Hu, Y. Liu, L. Zong, W. Zhang, R. Liu, Q. Xing, Z. Liu, Q. Yan, W. Li, H. Lei and X. Liu, Cell Death & Disease 14, 836 (2023) doi: 10.1038/s41419-023-06382-y

30 S.P. Langdon, S.S. Lawrie, F.G. Hay, M.M. Hawkes, A. McDonald, I.P. Hayward, D.J. Schol, J. Hilgers, R.C. Leonard and J.F. Smyth, Cancer Res 48, 6166–6172 (1988)

31 K. Katano, A. Kondo, R. Safaei, A. Holzer, G. Samimi, M. Mishima, Y.M. Kuo, M. Rochdi and S.B. Howell, Cancer Res 62, 6559–6565 (2002)

32 D.M. Provencher, H. Lounis, L. Champoux, M. Tétrault, E.N. Manderson, J.C. Wang, P. Eydoux, R. Savoie, P.N. Tonin and A.M. Mes-Masson, In Vitro Cellular & Developmental Biology - Animal 36, 357–361 (2000) doi: 10.1290/1071-2690(2000)036<0357:COFNEO>2.0.CO;2

33 P. Phadte, A. Bishnu, P. Dey, M. M, M. Mehrotra, P. Singh, S. Chakrabarty, R. Majumdar, B. Rekhi, M. Patra, A. De and P. Ray, Journal of Experimental & Clinical Cancer Research 43, 222 (2024) doi: 10.1186/s13046-024-03147-z

34 C.M. Beaufort, J.C. Helmijr, A.M. Piskorz, M. Hoogstraat, K. Ruigrok-Ritstier, N. Besselink, M. Murtaza, W.F. van Ĳcken, A.A. Heine, M. Smid, M.J. Koudijs, J.D. Brenton, E.M. Berns and J. Helleman, PLoS One 9, e103988 (2014) doi: 10.1371/journal.pone.0103988

35 S.C. Righetti, P. Perego, E. Corna, M.A. Pierotti and F. Zunino, Cell Growth Differ 10, 473–478 (1999)

36 S. Lederer, T.M.H. Dijkstra and T. Heskes, Frontiers in Pharmacology Volume 10 - 2019, (2019) doi: 10.3389/fphar.2019.01384

37 A. Ianevski, A.K. Giri and T. Aittokallio, Nucleic Acids Res 50, W739–w743 (2022) doi: 10.1093/nar/gkac382

38 B.N. Forgie, R. Prakash, A.A. Goyeneche and C.M. Telleria, Discov Oncol 15, 5 (2024) doi: 10.1007/s12672-023-00857-2

39 M.I. Love, W. Huber and S. Anders, Genome Biol 15, 550 (2014) doi: 10.1186/s13059-014-0550-8

40 G. Korotkevich, V. Sukhov, N. Budin, B. Shpak, M.N. Artyomov and A. Sergushichev, bioRxiv, 060012 (2021) doi: 10.1101/060012

41 A. Liberzon, C. Birger, H. Thorvaldsdóttir, M. Ghandi, J.P. Mesirov and P. Tamayo, Cell Syst 1, 417–425 (2015) doi: 10.1016/j.cels.2015.12.004

42 M. Kanehisa and S. Goto, Nucleic Acids Res 28, 27–30 (2000) doi: 10.1093/nar/28.1.27

43 S.B. Kovacs and E.A. Miao, Trends Cell Biol 27, 673–684 (2017) doi: 10.1016/j.tcb.2017.05.005

44 Z. Li, F. Mo, Y. Wang, W. Li, Y. Chen, J. Liu, T.J. Chen-Mayfield and Q. Hu, Nat Commun 13, 6321 (2022) doi: 10.1038/s41467-022-34036-8

45 C. Soldani, M.C. Lazzè, M.G. Bottone, G. Tognon, M. Biggiogera, C.E. Pellicciari and A.I. Scovassi, Exp Cell Res 269, 193–201 (2001) doi: 10.1006/excr.2001.5293

46 J.M. Weinberg, J.A. Davis, M. Abarzua and T. Rajan, J Clin Invest 80, 1446–1454 (1987) doi: 10.1172/jci113224

47 J.P. Borges, R.S.R. Sætra, A. Volchuk, M. Bugge, P. Devant, B. Sporsheim, B.R. Kilburn, C.L. Evavold, J.C. Kagan, N.M. Goldenberg, T.H. Flo and B.E. Steinberg, eLife 11, e78609 (2022) doi: 10.7554/eLife.78609

48 M. Raisova, A.M. Hossini, J. Eberle, C. Riebeling, T. Wieder, I. Sturm, P.T. Daniel, C.E. Orfanos and C.C. Geilen, J Invest Dermatol 117, 333–340 (2001) doi: 10.1046/j.0022-202x.2001.01409.x

49 T.G.O. Consortium, Nucleic Acids Research 54, D1779–D1792 (2026) doi: 10.1093/nar/gkaf1292

50 M. Ashburner, C.A. Ball, J.A. Blake, D. Botstein, H. Butler, J.M. Cherry, A.P. Davis, K. Dolinski, S.S. Dwight, J.T. Eppig, M.A. Harris, D.P. Hill, L. Issel-Tarver, A. Kasarskis, S. Lewis, J.C. Matese, J.E. Richardson, M. Ringwald, G.M. Rubin and G. Sherlock, Nature Genetics 25, 25–29 (2000) doi: 10.1038/75556

51 R. Skouta, S.J. Dixon, J. Wang, D.E. Dunn, M. Orman, K. Shimada, P.A. Rosenberg, D.C. Lo, J.M. Weinberg, A. Linkermann and B.R. Stockwell, J Am Chem Soc 136, 4551–4556 (2014) doi: 10.1021/ja411006a

52 A. Degterev, J. Hitomi, M. Germscheid, I.L. Ch’en, O. Korkina, X. Teng, D. Abbott, G.D. Cuny, C. Yuan, G. Wagner, S.M. Hedrick, S.A. Gerber, A. Lugovskoy and J. Yuan, Nat Chem Biol 4, 313–321 (2008) doi: 10.1038/nchembio.83

53 H. Hu, M. Tian, C. Ding and S. Yu, Front Immunol 9, 3083 (2018) doi: 10.3389/fimmu.2018.03083

54 K. Pakos-Zebrucka, I. Koryga, K. Mnich, M. Ljujic, A. Samali and A.M. Gorman, EMBO Rep 17, 1374–1395 (2016) doi: 10.15252/embr.201642195

55 H. Yoshida, T. Matsui, A. Yamamoto, T. Okada and K. Mori, Cell 107, 881–891 (2001) doi: 10.1016/s0092-8674(01)00611-0

56 J.M. Salvador, J.D. Brown-Clay and A.J. Fornace, Jr., Adv Exp Med Biol 793, 1–19 (2013) doi: 10.1007/978-1-4614-8289-5_1

57 R. Roufayel, K. Younes, A. Al-Sabi and N. Murshid, Life (Basel) 12, (2022) doi: 10.3390/life12020256

58 T. Kawase, H. Ichikawa, T. Ohta, N. Nozaki, F. Tashiro, R. Ohki and Y. Taya, Oncogene 27, 3797–3810 (2008) doi: 10.1038/onc.2008.32

59 T.J. Sayers, Cancer Immunol Immunother 60, 1173–1180 (2011) doi: 10.1007/s00262-011-1008-4

60 X. Ren, Z. Lin and W. Yuan, Front Pharmacol 13, 836614 (2022) doi: 10.3389/fphar.2022.836614

61 Z. Galehdar, P. Swan, B. Fuerth, S.M. Callaghan, D.S. Park and S.P. Cregan, J Neurosci 30, 16938–16948 (2010) doi: 10.1523/jneurosci.1598-10.2010

62 F. Meng, H. Li, Y. Wang, Z. Zheng and Y. Chen, Cell Investigation 1, 100046 (2025) doi: 10.1016/j.clnves.2025.100046

63 A. Gupta, C.R. Hunt, S. Chakraborty, R.K. Pandita, J. Yordy, D.B. Ramnarain, N. Horikoshi and T.K. Pandita, Radiat Res 181, 1–8 (2014) doi: 10.1667/rr13572.1

64 L.J. Mah, A. El-Osta and T.C. Karagiannis, Leukemia 24, 679–686 (2010) doi: 10.1038/leu.2010.6

65 C. Jiang, C. Shen, M. Ni, L. Huang, H. Hu, Q. Dai, H. Zhao and Z. Zhu, Genes & Diseases 11, 101063 (2024) doi: 10.1016/j.gendis.2023.06.032

66 Y. Xia, M. Sun, H. Huang and W.-L. Jin, Signal Transduction and Targeted Therapy 9, 92 (2024) doi: 10.1038/s41392-024-01808-1

67 L. Besse, A. Besse, S.C. Stolze, A. Sobh, E.A. Zaal, A.J. van der Ham, M. Ruiz, S. Phuyal, L. Büchler, M. Sathianathan, B.I. Florea, J. Borén, M. Ståhlman, J. Huber, A. Bolomsky, H. Ludwig, J.T. Hannich, A. Loguinov, B. Everts, C.R. Berkers, M. Pilon, H. Farhan, C.D. Vulpe, H.S. Overkleeft and C. Driessen, Cancer Research 81, 4581–4593 (2021) doi: 10.1158/0008-5472.Can-20-3323

68 M. Soprano, D. Sorriento, M.R. Rusciano, A.S. Maione, G. Limite, P. Forestieri, D. D’Angelo, M. D’Alessio, P. Campiglia, P. Formisano, G. Iaccarino, R. Bianco and M. Illario, PLOS ONE 11, e0155970 (2016) doi: 10.1371/journal.pone.0155970

69 X. Yang, F. Zheng, H. Xing, Q. Gao, W. Wei, Y. Lu, S. Wang, J. Zhou, W. Hu and D. Ma, J Cancer Res Clin Oncol 130, 423–428 (2004) doi: 10.1007/s00432-004-0556-9

70 C. Rogers, T. Fernandes-Alnemri, L. Mayes, D. Alnemri, G. Cingolani and E.S. Alnemri, Nature Communications 8, 14128 (2017) doi: 10.1038/ncomms14128

71 V. Zaitceva, G.S. Kopeina and B. Zhivotovsky, Cancers 13, 3671 (2021)

72 M. Nano, J.A. Mondo, J. Harwood, V. Balasanyan and D.J. Montell, Proceedings of the National Academy of Sciences 120, e2216531120 (2023) doi: doi:10.1073/pnas.2216531120

73 Y. Liu, L. Ouyang, S. Jiang, L. Liang, Y. Chen, C. Mao, Y. Jiang and L. Cong, Cancer Cell Int 24, 145 (2024) doi: 10.1186/s12935-024-03321-5

74 G. Eskander, S.G. Abdelhamid, S.A. Wahdan and S.M. Radwan, Cell Death Discovery 11, 56 (2025) doi: 10.1038/s41420-025-02328-9

75 S.L. Petersen, L. Wang, A. Yalcin-Chin, L. Li, M. Peyton, J. Minna, P. Harran and X. Wang, Cancer Cell 12, 445–456 (2007) doi: 10.1016/j.ccr.2007.08.029

76 O. Micheau, E. Solary, A. Hammann and M.-T. Dimanche-Boitrel, Journal of Biological Chemistry 274, 7987–7992 (1999) doi: 10.1074/jbc.274.12.7987

77 H. Yamaguchi and H.G. Wang, J Biol Chem 279, 45495–45502 (2004) doi: 10.1074/jbc.M406933200

78 K.M. Park, J.Y. Park, J. Pyo, S.Y. Lee and H.S. Kim, Molecules 27, (2022) doi: 10.3390/molecules27123804

79 D.R. Green, Cold Spring Harb Perspect Biol 14, (2022) doi: 10.1101/cshperspect.a041053

80 M. Lam, S.A. Marsters, A. Ashkenazi and P. Walter, eLife 9, e52291 (2020) doi: 10.7554/eLife.52291

81 H. Yamaguchi and H.-G. Wang, Journal of Biological Chemistry 279, 45495–45502 (2004) doi: 10.1074/jbc.M406933200

82 A.J. Cole, T. Dwight, A.J. Gill, K.-A. Dickson, Y. Zhu, A. Clarkson, G.B. Gard, J. Maidens, S. Valmadre, R. Clifton-Bligh and D.J. Marsh, Scientific Reports 6, 26191 (2016) doi: 10.1038/srep26191

83 A. Saleemuddin, A.K. Folkins, L. Garrett, J. Garber, M.G. Muto, C.P. Crum and S. Tworoger, Gynecol Oncol 111, 226–232 (2008) doi: 10.1016/j.ygyno.2008.07.018

84 A. Villunger, E.M. Michalak, L. Coultas, F. Müllauer, G. Böck, M.J. Ausserlechner, J.M. Adams and A. Strasser, Science 302, 1036–1038 (2003) doi: doi:10.1126/science.1090072

85 S. Oyadomari and M. Mori, Cell Death & Differentiation 11, 381–389 (2004) doi: 10.1038/sj.cdd.4401373

86 M. Wang, Z.M. Liu, X.C. Li, Y.T. Yao and Z.X. Yin, Journal of Chemotherapy 25, 162–169 (2013) doi: 10.1179/1973947812Y.0000000056

87 D.J. Peng, J. Wang, J.Y. Zhou and G.S. Wu, Biochem Biophys Res Commun 394, 600–605 (2010) doi: 10.1016/j.bbrc.2010.03.029

88 D. Zhang, H.-L. Piao, Y.-H. Li, Q. Qiu, D.-J. Li, M.-R. Du and B.K. Tsang, Experimental and Molecular Pathology 100, 506–513 (2016) doi: 10.1016/j.yexmp.2016.05.003

89 I. Gibbs-Seymour, E. Markiewicz, S. Bekker-Jensen, N. Mailand and C.J. Hutchison, Aging Cell 14, 162–169 (2015) doi: 10.1111/acel.12258

90 P. Knipscheer, M. Räschle, A. Smogorzewska, M. Enoiu, T.V. Ho, O.D. Schärer, S.J. Elledge and J.C. Walter, Science 326, 1698–1701 (2009) doi: 10.1126/science.1182372

91 A.J. Deans and S.C. West, Nat Rev Cancer 11, 467–480 (2011) doi: 10.1038/nrc3088

92 W. Sakai, E.M. Swisher, C. Jacquemont, K.V. Chandramohan, F.J. Couch, S.P. Langdon, K. Wurz, J. Higgins, E. Villegas and T. Taniguchi, Cancer Res 69, 6381–6386 (2009) doi: 10.1158/0008-5472.Can-09-1178

93 W. Sakai, E.M. Swisher, B.Y. Karlan, M.K. Agarwal, J. Higgins, C. Friedman, E. Villegas, C. Jacquemont, D.J. Farrugia, F.J. Couch, N. Urban and T. Taniguchi, Nature 451, 1116–1120 (2008) doi: 10.1038/nature06633

94 L. Zhang and X. Wang, Cell Death Discov 11, 444 (2025) doi: 10.1038/s41420-025-02761-w

