## Supplemental Figures for "The combination of nelfinavir and cisplatin drives lytic cell death through a caspase-8/caspase-3/GSDME axis in platinum-resistant ovarian cancer cells"

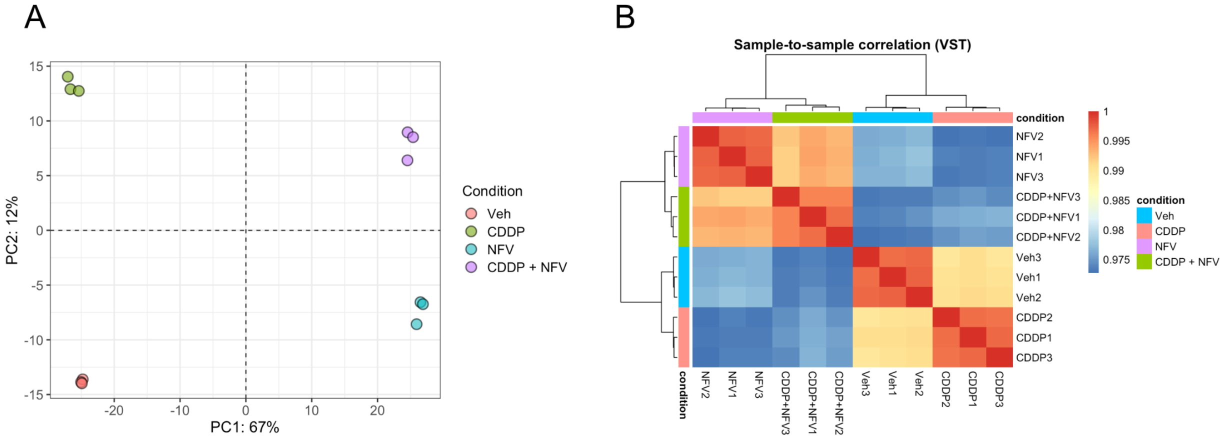


**Supplementary Figure S1. RNA-seq quality control demonstrates appropriate sample clustering and replicate consistency. A)** Principal component analysis (PCA) of variance-stabilized transcriptomic data from IGROV-1/CP cells treated with vehicle, 10 $\mu$M cisplatin (CDDP) for 3 h, 20 $\mu$M nelfinavir (NFV) for 24 h, or 10 $\mu$M CDDP for 3 h followed by 20 $\mu$M NFV for 24 h. Samples clustered according to treatment condition, with principal component 1 (PC1) and principal component 2 (PC2) accounting for 67% and 12% of the total variance, respectively.**(B)** Sample-to-sample correlation heatmap generated from variance-stabilized transcriptomic data showing strong within-group correlation and overall consistency among biological replicates across treatment conditions.

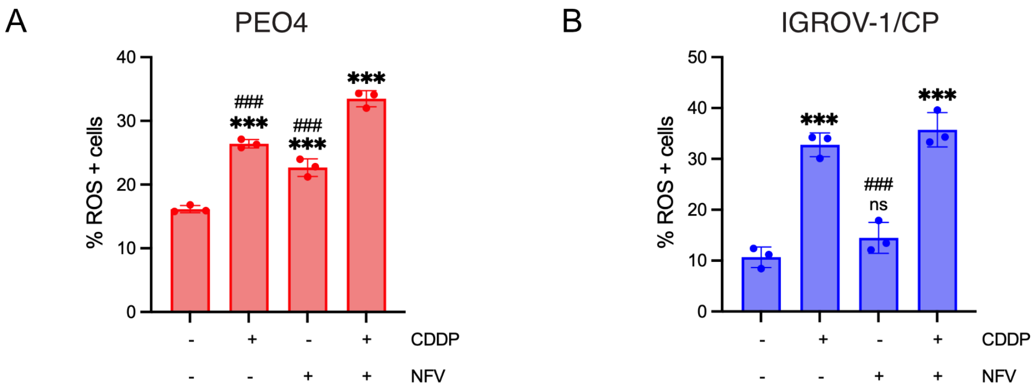


**Supplementary Figure S2. NFV/CDDP treatment induces reactive oxygen species in ovarian cancer cells. (A, B)** Reactive oxygen species (ROS) levels were measured in PEO4 (**A**) and IGROV-1/CP (**B**) cells following treatment with vehicle, 10 $\mu$M cisplatin (CDDP) for 3 h, 20 $\mu$M nelfinavir (NFV) for 72 h, or the combination, using the Muse Oxidative Stress Assay. In PEO4 cells, the NFV/CDDP combination increased the percentage of ROS-positive cells relative to vehicle and monotherapy controls. In IGROV-1/CP cells, ROS levels were also elevated following combination treatment, although the response was comparable to that observed with CDDP alone. Data are presented as mean ± SD from 2 independent experiments. Statistical analysis was performed using one-way ANOVA with Tukey's multiple comparisons test. *** $p < 0.001$ versus vehicle; ### $p < 0.001$ versus CDDP + NFV; ns, not significant.
